# Selective knockout of key CMV receptors in fetal cells blocks direct and endocytic pathways of entry in the guinea pig

**DOI:** 10.1101/2025.08.05.668711

**Authors:** Yushu Qin, K. Yeon Choi, Nadia El-Hamdi, Alistair McGregor

## Abstract

Cytomegalovirus is a leading cause of congenital disease and a vaccine is a high priority. Guinea pig with guinea pig cytomegalovirus (GPCMV) is the only small animal model for congenital CMV (cCMV). GPCMV encodes functionally essential viral entry glycoprotein complexes similar to HCMV, which are neutralizing antibody targets. As with HCMV, GPCMV has two pathways of cell entry (direct and endocytic). Common to both pathways and essential for infection is the fusogenic viral glycoprotein gB. Additional gH/gL-based complexes are necessary for receptor interaction and cell entry: gH/gL/gO trimer (direct); pentamer complex, PC (endocytic). Direct cell entry requires host PDGFRA receptor and viral trimer. An endocytic PC receptor has not been identified for GPCMV or any animal CMV. GPCMV endocytic entry was blocked by acidic flux inhibition but not direct cell entry, requiring knockout of PDGFRA. We hypothesized that cellular knockout of GPCMV direct and endocytic receptors would completely block infection. Two PC receptor candidates, guinea pig NRP2 and CD147, present on all established guinea pig cell lines were evaluated. Results demonstrated that NRP2 interacted with PC unlike CD147 in immunoprecipitation assays. Double knockout of PDGFRA and NRP2 completely blocked GPCMV but had no impact on control HSV-1 infection. In contrast, CD147/PDGFRA double knockout had limited inhibition of GPCMV and no impact on HSV-1. Ectopic expression of cell receptors restored infection to normal levels on knockout cell lines. Overall, results demonstrate GPCMV conservation with HCMV for key receptors and cell entry pathways enhancing the translational importance of this model.

**Importance:** Congenital CMV is a leading cause of hearing loss and cognitive impairment in newborns and a vaccine is a high priority. Species-specificity of HCMV requires animal model studies to utilize species-specific virus. The guinea is the only small animal model for cCMV and GPCMV encodes functional HCMV homolog glycoprotein complexes for cell entry via direct or endocytic pathways. The gB glycoprotein is required for infection of all cell types but a gB vaccine fails to fully protect against cCMV. GPCMV encodes a functional PC required for infection of different cell types via endocytic pathway. The PC has emerged as an important vaccine antibody target but PC-based cell entry is only partially understood and poorly characterized for GPCMV. Identifying GPCMV cell entry receptors is critical to the understanding of virus tropism and disease in this model. Correlation with HCMV improves translational impact of a GPCMV vaccine and antiviral cCMV intervention.

## Introduction

Human cytomegalovirus (HCMV), a common betaherpesvirus, establishes a lifelong persistent infection with severest disease associated with immunosuppressed populations including transplant and AIDS patients (1). Additionally, HCMV has the ability to cross the placenta and cause congenital cytomegalovirus (cCMV) infection with symptomatic disease resulting in vision and cognitive impairment as well as sensorineural hearing loss (SNHL) in newborns and is associated with autism (2, 3). In Europe and the US, cCMV occurs in approximately 0.5-1.2% of newborn babies with up to 30% of hearing loss in children attributed to cCMV (4–6). Globally over a million babies are born each year with cCMV (2) and a vaccine is considered a high priority (7). HCMV species specificity complicates studies in preclinical animal model, which requires the use of species-specific animal CMV. The guinea pig is the only small animal model for cCMV with guinea pig cytomegalovirus (GPCMV) causing disease in newborn pups similar to that in humans including SNHL (8–11). The guinea pig has a hemomonochorial placenta structure similar to humans and pregnancy studies evaluated in trimesters with prenatal pup neuro anatomical development occurring almost completely in utero (12–14). Various intervention strategies against cCMV have been evaluated in this model (11, 15).

HCMV viral glycoproteins form specific complexes on the virion that are required for direct cell entry (gB, gM/gN, and gH/gL/gO), and function as neutralizing-antibody targets (16–21) which enable infection of cells by a direct pathway of entry. Clinical strains of HCMV also encode an additional gH/gL-based pentamer complex, or PC (gH/gL/UL128/UL130/UL131) which in association with gB enables cell entry by an endocytic pathway for predominantly infection of epithelial, endothelial and myeloid cells which lack or express low levels of PDGFRA (22, 23). Guinea pig CMV (GPCMV) encodes homolog glycoprotein complexes (gB, gH/gL/gO, gM/gN) essential for GPCMV infection which also act as neutralizing target antigens (11, 24–26). GPCMV also encodes a homolog PC (gH/gL/GP129/GP131/GP133) necessary for endocytic pathway of infection in conjunction with gB (25). Cellular platelet-derived growth factor receptor alpha (PDGFRA), present mainly on fibroblasts, has been identified as the cell receptor necessary for HCMV direct fusion pathway of cell entry (27). Guinea pig PDGFRA is required for GPCMV direct infection of fibroblast cells with the receptor interacting with viral gH/gL/gO trimer but viral cell entry also required gB (26, 28). Various guinea pig cell lines have recently been established with GPCMV epithelial and endothelial infection requiring PC for endocytic entry as they lack PDGFRA (26, 29, 30). However, PDGFRA can act as a universal receptor for GPCMV direct cell entry in all cell types by ectopic expression (29, 30).

In both HCMV and GPCMV, the gB glycoprotein is an immunodominant neutralizing target and essential for infection of all cell types as the fusogenic viral glycoprotein (24, 25, 31–36). Consequently, the gB complex is considered an essential component of any vaccine against congenital CMV. However, a gB (HCMV) subunit vaccine, despite evoking high antibody titers, exhibited only approximately 50% efficacy in phase II clinical trials (37, 38). GPCMV gB subunit vaccine strategies in the guinea pig model fail to fully protect against congenital CMV and lack cross strain protection (39) exhibiting efficacy similar to that in human gB vaccine trials (11). In both HCMV and GPCMV, gB antibodies are less effective at neutralizing virus infection of non-fibroblasts than fibroblasts, which might be a basis for reduced vaccine efficacy (35, 40–42). Other target antigens are being explored as vaccine candidates including the PC as a neutralizing antibody target (23, 43, 44). Several human receptors candidates have been identified for PC-dependent cell entry of HCMV, including neuropilin-2 (NRP2), OR14I1, CD147, CD46, and thrombomodulin (ThBD) (45–50). The process of PC-dependent cell entry is only partially understood, and receptors are likely to vary between cell types. For example, a specific PC receptor for trophoblast has not been identified. GPCMV PC is necessary for infection of cells lacking PDGFRA including epithelial, endothelial, and trophoblast cells and macrophage (11, 25, 29, 30, 51, 52). The PC is therefore highly important for GPCMV dissemination and congenital infection (25, 29, 30, 51). Inclusion of a PC in a vaccine strategy against GPCMV dramatically improves neutralizing-antibody titers and contributes to enhancement of protection against cCMV (26, 53). Given the importance of GPCMV PC to virus infection and the development of an effective CMV vaccine, it was critical to gain a better understanding of viral infection by identifying candidate PC cellular receptors for GPCMV.

In prior studies, CRISPR/Cas9-based knockout (KO) strategy of PDGFRA on guinea pig fetal lung GPL fibroblast cells rendered them refractile to GPCMV(PC-) infection by blocking the direct route of cell entry. However, guinea pig PDGFRA KO cells remained susceptible to infection by GPCMV(PC+) virus via endocytic pathway with similar growth kinetics as infection of wild type GPL cells (26, 28). We hypothesized that targeted knockout of receptors for both direct and endocytic pathways would completely block GPCMV infection. Additionally, PC/receptor interaction could be confirmed by transient expression immunoprecipitation assay. Initial, screening of various established guinea pig cell lines, which were capable of PC-dependent endocytic infection identified two candidates expressed in all cell lines. Guinea pig NRP2 and CD147, also known as basigin (BSG), which were expressed in various cell lines including, fibroblast, epithelial and endothelial cells. CRISPR/Cas9-based double knockout (DKO) of PDGFRA and NRP2 on GPL cells resulted in the complete blocking of GPCMV(PC+) infection. Additionally, a PDGFRA KO GPL cell line positive for NRP2, demonstrated that PC-dependent pathway of entry, which could be inhibited by preventing the acidic endocytic entry flux by chemical treatment of cells with Bafilomycin A (BaflA) similar to epithelial cells. A separate CRISPR-based DKO of PDGFRA and CD147 in GPL cells, resulted in reduction of GPCMV(PC+) infection but not complete inhibition. Control HSV-1 infection of GPL or various KO cell lines was unaffected by specific receptor knockout with inhibition directed specifically to GPCMV. Transient expression of PC and candidate receptors in immunoprecipitation (IP) assays demonstrated specific interaction of GPCMV PC with guinea pig NRP2 but not CD147. Viral trimer (gH/gL/gO) IP assays lacked interaction with CD147 or NRP2 but did demonstrate interaction with PDGFRA. Ectopic expression of PDGFRA, NRP2 or CD147 on respective KO or DKO cell lines restored GPCMV infection to near normal levels. Overall, results demonstrate that GPCMV utilizes NRP2 as a PC-based entry receptor similar to HCMV but that CD147 has potential to impact PC-based cell entry indirectly.

## Materials & Methods

### Cells, viruses, oligonucleotides and genes

GPCMV (strain 22122, ATCC VR682) was propagated on various cell lines. Cell lines used in studies included: Guinea pig fibroblast lung cells (GPL; ATCC CCL 158), renal epithelial cells (REPI); guinea pig amniotic sac (GPASE); guinea pig umbilical cord endothelial cells (GPUVEC); guinea pig trophoblasts (TEPI) (25, 29, 30, 51). GPCMV BAC derived virus (PC+/PC-/GFP+/-) were generate as previously described (25). Recombinant defective adenovirus (Ad5) vectors (E1 and E3 deleted) were previously described encoding individual components of the PC (gH, gL, GP129, GP131, GP133) and additional recombinant Ad vectors encoded gO, control GFP, or mCherry reporter gene (35, 54). All ORFs were under HCMV IE enhancer control with 3’ SV40 polyA sequence. High titer CsCl gradient purified recombinant defective adenovirus virus stocks (10^12^ TDU/ml) were generated by Welgen Inc. (MA). All oligonucleotides were synthesized by Sigma-Genosys (The Woodlands, TX). Synthetic codon optimized genes were developed for: [1] guinea pig NRP2 and [2] guinea pig CD147 (Genscript). Guinea pig ORFs were derived from guinea pig NCBI genome sequence (Cavia porcellus annotation release 103 GCF_000151735.1): NRP2 (XM_013157537); and CD147 (BSG) (XM_004999221). Both NRP2 and CD147 ORFs additionally incorporated a C-terminal FLAG epitope tag. The ORFs were cloned into pcDNA3.1(+) vector (Invitrogen) under HCMV IE enhancer promoter control to enable transient expression in plasmid transfected cells with protein expression verified by estern blot.

### Western blot assays

Western blots were carried out on cell lysates separated by 4-20% SDS-PAGE under denaturing conditions and performed as previously described (24, 25, 35). For western blots, anti-epitope tag primary antibodies were used at 1/1000: FLAG (Novus Biological); GFP (Santa Cruz Biotechnology); His (Bethyl): MYC-c (Novus Biologicals); and mCherry (Clontech Laboratories) (25, 26). Secondary antibodies: anti-mouse or anti-rabbit IgG HRP-linked secondary antibodies for western blot (Cell Signaling) were used at 1/2000 (26). FLAG epitope was used to detect CD147, NRP2, GP133 and gO protein expression in transduced/transfected cell monolayers. Additionally, species cross-reactive antibodies were used for the detection of NRP2 (R&D Systems), CD147 (Antibodies Online), and PDGFRA (R&D Systems).

### Immunoprecipitation assays

Immunoprecipitation (IP) assays were carried out on plasmid transfected or recombinant Ad transduced guinea pig cells using commercial RFP-trap (ChromoTek) following manufacturer’s protocols with inclusion of protease inhibitor cocktail (Pierce) in cell lysates. Samples were subsequently analyzed by SDS-PAGE (4–20% gradient gel) and western blot using specific anti-epitope tag antibodies: FLAG (Novus Biological); GFP (Santa Cruz Biotechnology); His (Bethyl): MYC (Novus Biologicals); and mCherry (Clontech Laboratories). Appropriate secondary anti-mouse or anti-rabbit HRP conjugate (Cell Signaling Technology) were also used following standard western blot protocol as previously described (25).

### Bafilomycin A inhibition of virus entry by pH-dependent endocytosis

Bafilomycin A1 (BaflA) inhibition of GPCMV infection was carried out following a previously described protocol (25). GPL fibroblast cells in six well plates were pretreated with complete media containing either 0 or 100 nM BaflA (Sigma) for 1 hr at 37°C followed by GPCMV(PC+) GFP-tagged virus infection (MOI = 0.5 pfu/cell) for 1hr at 37 °C. All further incubations were performed at the same concentration as pretreatment in complete media. Cells were fixed at 24 hr post infection with 4% PFA (15 min) and subsequently overlayed with PBS and stored at 4° C as sealed plates. Infected cells were identified by GFP fluorescent microscopy. Infection was set up in triplicate. Counts were made of GFP positive cells in random fields. Statistical analysis was performed using student T-test on the percent of cells infected in thirty random fields of view, each contained ∼100 cell nuclei, for each condition. The number of treated cells infected was represented as a percentage of the number of infected untreated cells.

### Flow cytometry and detection of cell surface expression of PDGFRA and NRP2

Flow cytometry antibody detection of PDGFRA (R&D Systems) and NRP2 (R&D Systems) cell surface expression was carried following manufacturer’s protocol. Surface protein detection for endogenous expression of PDGFRA and NRP2 were tested on wild type and both DKO (NRP2/PDGFRA & CD147/PDGFRA) GPL cells. Cells were grown on 6 well plates and samples were evaluated in duplicate. GPL cells (wildtype or DKOs) were labeled with primary antibody (anti-PDGFRA or anti-NRP2) 0.25µg/10^6^ cells in 100 μl of flow buffer (1X PBS + 2% FBS). Cells were washed with flow buffer 2X then labeled with 100 μl secondary antibody (donkey anti-goat AF488 or donkey anti-goat AF647, Abcam) depending on cell line used, diluted 1/2000 in flow buffer. Unlabeled and secondary antibody only GPL cells were included as unstained antibody controls. All incubations occurred for 30 min on ice in the dark. Flow cytometry was carried out on a Fortessa X-20 (BD Life Sciences) and data analyzed by FlowJo software (V10).

### CRISPR/Cas9 mutagenesis knockout strategy

*Guinea pig PDGFRA gene knockout.* Knockout of PDGFRA in GPL cells was carried out as previously described (26). The sequence of the guinea pig PDGFRA (gpPDGFRA) gene (NCBI gene accession number 100726209) was based on the guinea pig NCBI genome sequence (Cavia porcellus annotation release 103 GCF_000151735.1), and predicted introns and exons were additionally identified via use of the genome with Ensembl accession number ENSCPOG00000011782. Exon 2 of gpPDGFRA was targeted for mutagenesis by use of the CRISPR/Cas9 strategy. Exon 2-specific guide RNA (gRNA) was designed via an online program (www.rgenome.net) to avoid off-target sites. DNA sequences were cloned under US6 promoter control in three separate gRNA expression plasmids (pCas-Guide-EF1a-GFP; OriGene): pR1 (5′-GGTGTGGGCCGCCGAGGCGT-3′), pR2 (5′-TCTGGGAGAGTTCCCCGACG-3′), and pR3 (5′-CGTTTCTGATGTCCACGTCG-3′). GPL cells in 6-well plates were transduced with defective lentivirus expressing Cas9 under HCMVIE enhancer control (Origene) to enable expression of Cas9 and subsequently transfected with the gRNA expression plasmids (pR1, pR2, and pR3). At 2 days post-transfection, the cells were reseeded in 96-well plates by limiting dilution to generate individual cell lines. PDGFRA gene knockout cell lines were screened by exon 2 PCR sequencing of extracted genomic DNA. Genomic extraction was carried out with a DNeasy extraction kit (Qiagen), and PCR was performed using primers Fpd (5′-CTGAGCCTAATCTGCTGCCAGCTTTCG-3′) and Rpd (5′-CGGCACGGTAGATGTAGATATGC-3′). Western blotting of total cell lysate for specific modified cell lines verified the loss of gpPDGFRA expression. Goat anti-mouse PDGFRA antibody (R&D Systems) was predicted to react with gpPDGFRA based on 100% conservation of the target sequence, and this was verified by western blot analysis of cells transfected with a transient expression plasmid encoding a synthetic full-length gpPDGFRA with a C-terminal FLAG epitope tag (GenScript). The PCR products of the wild-type GPL and PDGFRA mutant knockout cell lines were cloned into the TA cloning vector (Invitrogen) and sequenced as previously described (28). Alignment of the PDGFRA exon 2 DNA sequence with wild-type GPL and GPKO cells was previously shown (26) and modification to PDGFRA in exon 2 of double knockout cell lines (NRP2/PDGFRA and CD147/PDGFRA) was as previously described (26). Double gene knockout strategy required the use of additional gRNA plasmids targeting either NRP2 or CD147.

*NRP2 and CD147 gRNA.* Guinea pig NRP2 (XM_013157537) and CD147 (BSG) (XM_004999221) gene sequence were derived from guinea pig NCBI genome sequence (Cavia porcellus annotation release 103 GCF_000151735.1) as described for PDGFRA. Predicted introns and exons were additionally identified via use of the genome analysis with Ensembl with accession number ENSCPOG00000004390 for NRP2 and ENSCPOG00000023530 for BSG. Exon specific gRNAs were designed via online program www.rgenome.net/be-designer/ to avoid off-target sites. DNA sequences were cloned under US6 promoter control in three separate gRNA expression plasmids (pCas-Guide, OriGene) for each target gene. NRP2 gRNA plasmids: nR1. 5’ATGGACCATCGAATCTCCG3’; nR2. 5’GATGATCTCCATCTTGGGTT3’; nR3. 5’AGGGTCATGCTCCAGGTCAA3’; nR4. 5’ATGGGGACAGCGAGTCCG3’. CD147 gRNA plasmids: bR1. 5’ACAGTGATGTCGCTGTTACG3’; bR2. 5’CGTGGGCTCGCGGGTGCACA3’; bR3. 5’ATCACTGTGGTTCAGGACGC3’. *Generation of double knockout GPL cell lines (PDGFRA/NRP2 DKO and PDGFRA/CD147).* Generation of double knockout cell lines followed the same protocol described for single PDGFRA knockout with Cas9 positive GPL cell transfected with appropriate sets of gRNA expression plasmids (either PDGFFRA+NRP2 gRNA or PDGFRA+CD147). Cells were seeded by limiting dilution and cell lines were subsequently screened by PCR exon sequencing for DNA modification and western blot for loss of target protein expression to confirm double gene knockout using PDGFRA primers described above as well as specific primers for NRP2 or CD147. NRP2 PCR primers Fnrp2 (5’CATCCAGGTATGACTTCATCG3’) and Rnrp2 (5’CTCTTCCACCTGTCACGC3’). CD147 PCR primers Fbsg (5’GTGACCACCATGGACAG3’) and Rbsg (5’TCCTCCTTCAGCACCTTG’3’). Confirmed double knockout cell lines were designated as DKO NRP2/PDGFRA GPL cells for knockout of both NRP2 and PDGFRA or DKO CD147/PDGFRA GPL cells for knockout of both CD147 and PDGFRA.

### BLAST protein alignment

Alignment and predicted identity for candidate receptors (NRP2 and CD147) between guinea and human counterparts was carried out with predicted amino acid sequence of guinea pig and human proteins based from NCBI data base with alignment carried out with NCBI BLASTp program (https://blast.ncbi.nlm.nih.gov/Blast.cgi). NCBI accession numbers for predicted proteins: Guinea pig NRP2 (XP_013012991); Guinea pig CD147 (BSG) (XP_004999278); Human NRP2 (NP_957718); Human CD147 (BSG or EMMPRIN) (BAB88938.1).

### Statistical analysis

All statistical analyses were conducted with GraphPad Prism (version 7) software. Fisher’s exact test, Student’s t test (unpaired) with significance taken as a *p* value of <0.05 or as specified in the figure legends.

## Results

### Evaluation of candidate receptor NRP2 expression and interaction with GPCMV PC

In HCMV, NRP2 is the most commonly associated cellular receptor for PC based virus infection. The guinea pig genome encodes a homolog NRP2 protein with high identity (93%) to human NRP2 based on BLAST protein analysis of their respective predicted amino acid sequence between human and guinea pig NRP2 proteins. Based on the predicted ORF a codon optimized guinea pig NRP2 synthetic gene was generated (Genscript) encoding a C-terminal FLAG epitope tag and cloned into pcDNA3.1(+) under HCMV IE enhancer control for transient plasmid expression studies (pDNAgpNRP2). A protein of approximately 115 kDa was detected by anti-FLAG western blot of plasmid transfected GPL cell lysate which was similar to the size of human NRP2 and predicted guinea pig protein size (Fig 1(i)). All available previously characterized guinea pig cell lines were evaluated for expression of NRP2 protein by western blot of total cell lysates using a cross reactive anti-NRP2 antibody (R&D Systems) which also detected synthetic NRP2 protein expression (pDNAgpNRP2) by western blot analysis verifying cross reactivity of the antibody (Fig 1(iii)). Guinea pig cell lines tested for NRP2 expression included: amniotic sac (GPASE); trophoblast (TEPI); umbilical cord endothelial (GPUVEC); renal epithelial (REPI); fetal fibroblast lung (GPL) cells; and GPL cells + NRP2 plasmid (25, 29, 30, 51). Western blot analysis identified a 115 kDa protein on all cell lysates evaluated and indicated NRP2 protein was expressed on various guinea pig cell types (Fig1(iii)). However, the level of NRP2 expression varied between different cell lines based on lane loading control (β-actin) and strongly expressed on GPL cells, Fig 1(iii).

**Figure 1.**
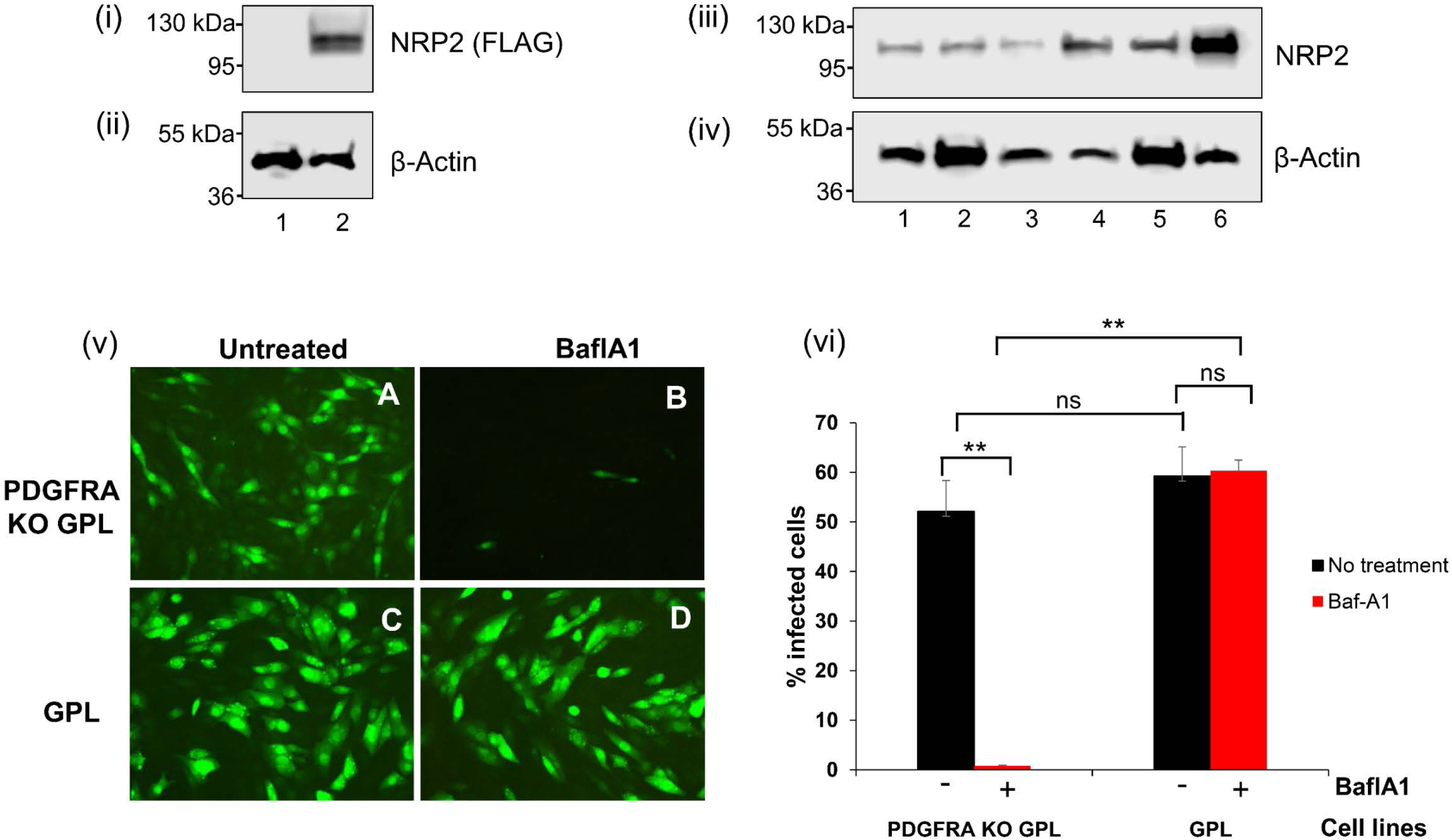
Detection of NRP2 protein expression in guinea pig cells by western blot and inhibition of GPCMV endocytic cell entry by pretreatment of cells with acidic flux inhibitor BaflA. (i)-(iv) Western blot detection of NRP2. (i) Detection of transient plasmid FLAG-tagged NRP2 synthetic gene protein expression on GPL cells by western blot of cell lysate. Lanes: 1, Control GPL cells; 2, NRP2 plasmid transfected cells. (ii) β-actin western lane loading control for (i). (iii) Western blot of endogenous NRP2 expression on various guinea pig cell lines with anti-NRP2 antibody. Lanes: 1, GPASE; 2, TEPI; 3, GPUVEC; 4, REPI; 5, GPL; 6, GPL+ NRP2 plasmid. (iii) β-actin western lane loading control for (iii). (v)-(vi) Inhibition of endocytic GPCMV infection by BaflA. GPL or PDGFRA KO GPL cells were untreated or pretreated with 100 nM BaflA for 1 hr prior to GPCMV(GFP+PC+) infection (MOI = 0.5 pfu/cell). (v) GFP+ virus infection in presence or absence of BaflA. Images of random cell fields for GFP positive cells. Untreated cells infected with virus (images A & C); pretreated with BaflA (images B & D). GFP+ cells imaged at 10X magnification at 36 hpi. (vi) Percentage of GPCMV infected cells (GPL or PDGFRA KO GPL) either BaflA treated or untreated with 10 random fields counted per condition based on 3 independent experiments and average values shown. Statistical analysis was performed with student t-test (\*\**p* < 0.01; ns = not significant).

Previously, we demonstrated that GPL cells expressed PDGFRA which enabled cell entry by PC-independent direct pathway (28). CRISPR knockout of PDGFRA (PDGFRA KO GPL cells) prevented GPCMV(PC-) direct cell entry but not GPCMV(PC+) endocytic cell entry with infection rate similar to that of wild type GPL cells (26, 28). In other guinea pig cell lines, endocytic PC-based infection was dependent upon an acidic flux similar to HCMV, which could be blocked by bafilomycin A (BaflA) (25, 51). In order to assess whether endocytic infection of PDGFRA KO fibroblast cells was also dependent upon a pH flux, GPCMV(PC+) infection was evaluated on GPL cells and previously established PDGFRA KO GPL fibroblasts (26, 28). GPCMV(PC+) infection (0.5 pfu/cell) was carried out in triplicate in six-well plates in the presence or absence of BaflA pretreatment as previously described using GFP-tagged GPCMV(PC+) virus (25). At 24 hr post infection (hpi) cells were fixed and GFP positive cells counted in replicate fields with representative untreated and treated cells shown in Fig 1(v). Overall results shown as average percentage of infected cells for GPL PDGFRA KO or GPL cells under different conditions (Fig 1(vi)) demonstrated that virus infection could be inhibited by BaflA treatment of PDGFRA KO cells unlike wild type GPL cells based on detection of GFP positive cells. It was concluded that PC endocytic infection of GPL cells was similar to that of GPCMV (PC+) infection (25, 51) of PDGFRA negative non-fibroblast cells requiring an acidic flux for endocytic cell entry and GPL cells were suitable for further receptor candidate KO studies.

Prior to attempted receptor candidate knockout in guinea pig cells, it was important to demonstrate interaction of viral PC with NRP2 protein. Transient protein expression immunoprecipitation (IP) assays were used to evaluate PC interaction with both endogenously expressed and plasmid-based NRP2 protein. A RFP/mCherry trap (ChromoTek) strategy was employed to immunoprecipitate C-terminal mcherry-tagged gL protein and interacting components of the PC as well as potential receptor NRP2 protein following a previously described protocol for GPCMV PC IP assay (25). GPL cells were transduced with recombinant defective adenovirus (Ad) vectors encoding individual components of the PC with individual protein carrying unique C-terminal epitope tags (Fig 2(i)): gH (GFP); gL (mCherry); GP129 (MYC); GP131 (His); GP133 (FLAG). Cells were initially transfected with FLAG-tagged NRP2 expression plasmid (pDNAgpNRP2) prior to transduction with Ad vectors and subsequent IP (25). Cell lysate and IP samples were analyzed by SDS-PAGE and western blot for detection of individual PC components and NRP2 protein. Results (Fig 2(ii)) demonstrated that for the control gH/gL subcomplex there was no detectable interaction with endogenous or plasmid expressed NRP2 protein in IP assays. In contrast, expression of the full PC in an IP assay successfully pulled down all components of the PC as well as NRP2 protein, both plasmid expressed and endogenous (Fig 2(iii)). This indicated that GPCMV PC was required for interaction with NRP2 and suggesting that NRP2 was a candidate entry receptor.

**Figure 2.**
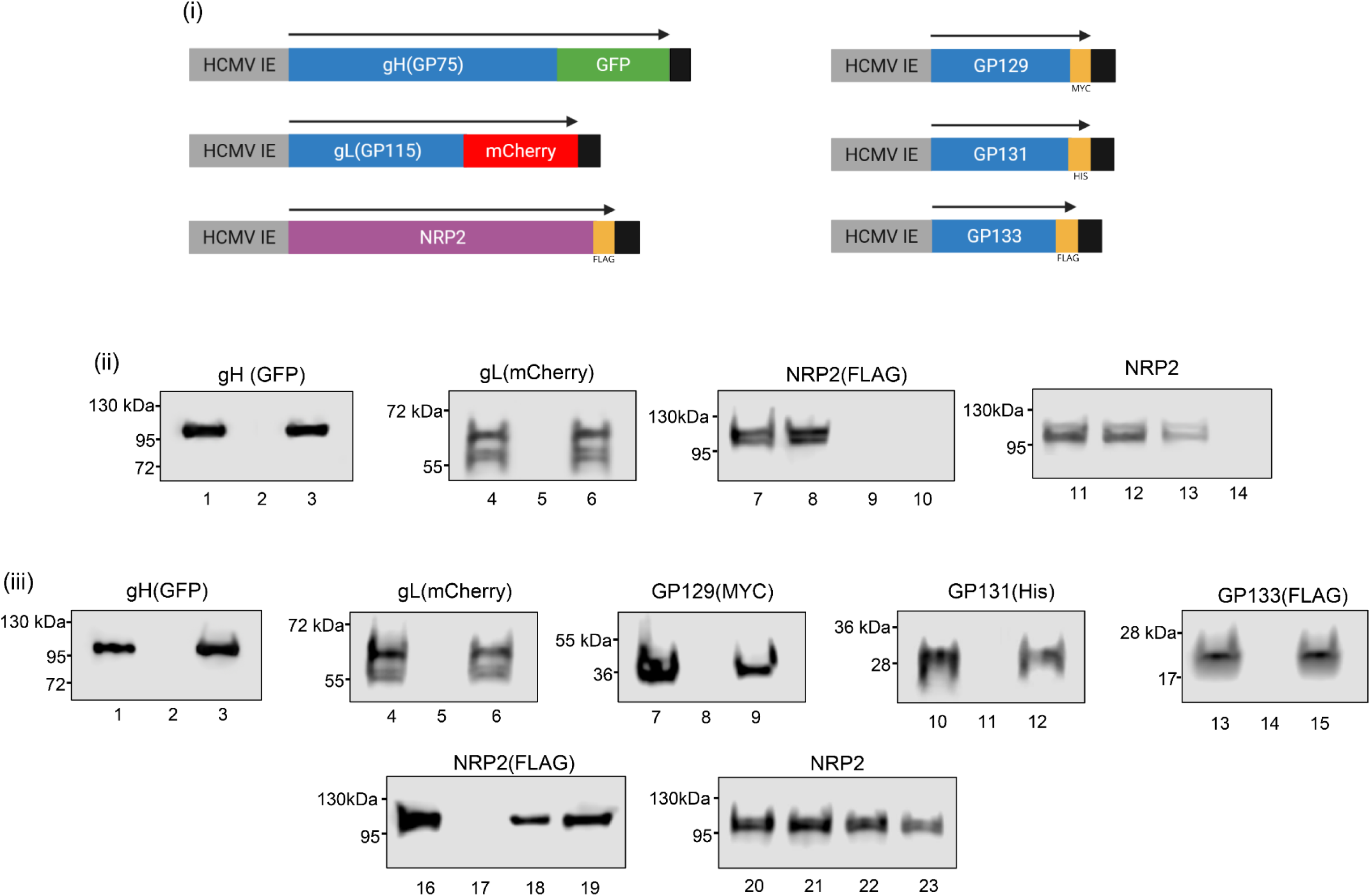
Immunoprecipitation (IP) of PC and candidate receptor NRP2. (i) Diagram of specific expression cassettes used for individual PC components and NRP2. (ii) Control IP assay with gHGFP, gLmcherry and NRP2. Cells transduced with Ad vectors (10 TDU/cell/virus) encoding gH(α-GFP) and gL(α-mCherry) and NRP2(α-FLAG) plasmid transfection followed by RFP IP and western blot for proteins. Lanes: 1-3 (GFP); 4-6 (mCherry); 7-10 NRP2 (FLAG); 11-14 α-NRP2 (endogenous). Lanes 1, 4, 7, 11 (total cell lysate). Lanes 3, 6, 10, 14 (IP). Lanes 2, 5, 9, 12 (mock control lysate). Lanes 8 and 13 IP flow through. Protein detected with appropriate secondary antibody-HRP conjugate as described in materials and methods. (iii) Western blot of PC IP assay. GPL cells transduced with five recombinant Ad vectors encoding individual PC components (10 TDU/virus/cell) and NRP2 plasmid transfection. Samples analyzed as total cell lysate or processed for immunoprecipitation (IP) assay. IP carried out with RFP-trap as previously described (25). IP lysate analyzed by western blot using respective primary antibodies for individual PC components or NRP2 (plasmid or endogenous): gH (α-GFP); gL (α-mCherry); GP129 (α-MYC); GP131 (α-His); GP133 (α-FLAG); endogenous NRP2 (α-NRP2); plasmid NRP2 (α-FLAG). Western blot lanes: 1-3 (GFP); 4-6 (mCherry); 7-9 (MYC); 10-12 (His); 13-15 GP133 (FLAG); 16-19 NRP2 (FLAG); 20-23 (NRP2 for endogenous). Lanes 1, 4, 7, 10, 13, 16, 20 (total cell lysate). Lanes 3, 6, 9, 12, 15, 19, 23 (IP). Lanes 2, 5, 8, 11, 14, 17, 21 (mock control lysate). Lanes 18 and 22 IP flow through.

### Blocking GPCMV infection by double knockout of PDGFRA and NRP2 receptors

CRISPR/Cas9 gene knockout was carried out on GPL cells as previously described (26, 28) utilizing three gRNA targeting PDGFRA (exon 2) and four gRNA targeting NRP2 (exon 2/3). Resulting double knockout cell line (NRP2/PDGFRA DKO) was characterized to confirm ablation of both PDGFRA and NRP2 protein expression. Cell lysates of GPL and DKO cell lines were used to evaluate PDGFRA and NRP2 expression by western blot assays (Fig 3(i-iii) to confirm absence of detectable protein in the DKO cell line. Specific alteration of PDGFRA and NRP2 targeted exon gene sequence was also confirmed by PCR sequencing (data not shown). Next, separate one-step growth curves (MOI 1pfu/cell) were performed on confluent monolayers of GPL and DKO cells in six well plates with GPCMV(PC+) virus (strain 22122) stock generated on renal epithelial cells. Results (Fig 3 (iv)) demonstrated failure of the virus to grow on DKO cells compared to normal growth kinetics on GPL cells. Virus also replicated normally on PDGFRA KO cell line (26) (data not shown). Comparative images of DKO and GPL cells either infected or control mock infected cells at 3 days post infection (Fig 3(v)) demonstrated that viral cpe from infection was associated only with infected GPL cells. Clinical GPCMV strain TAMYC which grew on GPL cells also failed to grow on NRP2/PDGFRA DKO cells (data not shown) (39). Transient ectopic expression of PDGFRA and NRP2 following previously described protocol (28) restored GPCMV growth to near wild type levels on DKO cells (Figure 3(vi)). Additionally, flow cytometry was performed (Fig 4) to confirm surface expression of PDGFRA and NRP2 on GPL cells and absence of expression on DKO cell line which was predicted from lack of detectable protein by western blot (Fig 3(i)). Overall, it was concluded that NRP2 was capable of acting as receptor for PC-dependent GPCMV cell entry and knockout of both PDGFRA and NRP2 blocked both pathways of cell entry for GPCMV.

**Figure 3.**
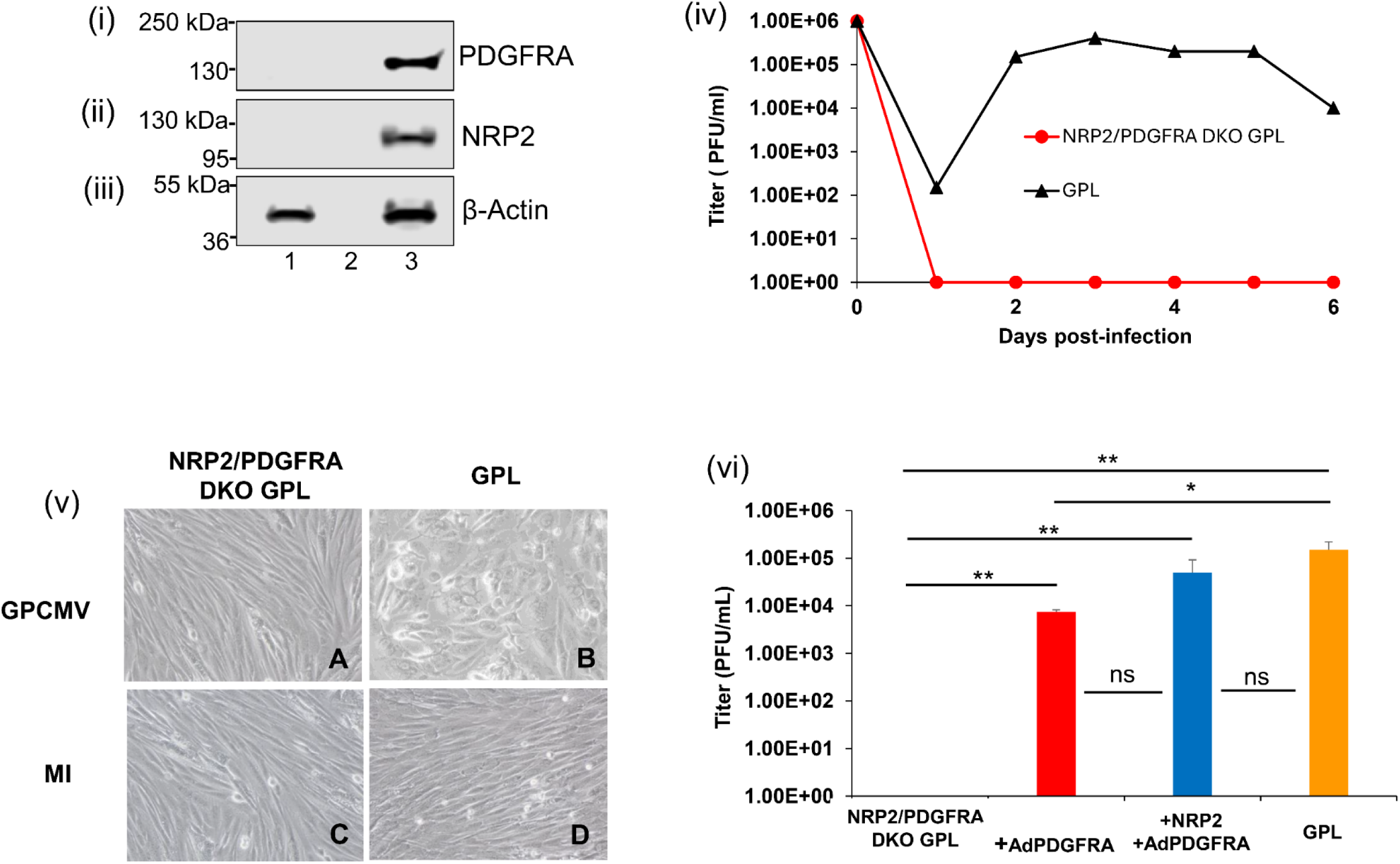
Characterization of NRP2/PDGFRA DKO cells and evaluation of GPCMV infection. (i)-(iii) Receptor protein western blot analysis of cell lysate. (i) Western blot for PDFGRA. (ii) Western blot for NRP2. (iii) Western blot for lane loading control β-actin. Lanes: 1, DKO cells; 2, loading dye; 3, GPL cells. (iv) Comparative growth curve of GPCMV(PC+) on GPL (black triangle) and DKO cells (red circle). Virus infection MOI 1 pfu/cell with evaluation of days 1-6 post infection. Results shown are average viral titers for experiment carried out in triplicate on 6 well plates with infection on confluent cell monolayers. (v) Bright field images of DKO (A & C) or GPL cells (B & D) either infected (A & B) or mock infected (C & D). Images represent random fields taken at 4 days post infection. Input virus MOI 1 pfu/cell. (vi) Comparative GPCMV(PC+) growth (MOI 1 pfu/cell) on DKO (black) or GPL (orange) cells or DKO cells ectopically expressing receptors PDGFRA (red) or NRP2+PDGFRA (blue). Results shown are average virus titers at 3 days post infection from experiments performed in triplicate (\*\**p* < 0.01; \**p* < 0.05; ns = not significant).

**Figure 4.**
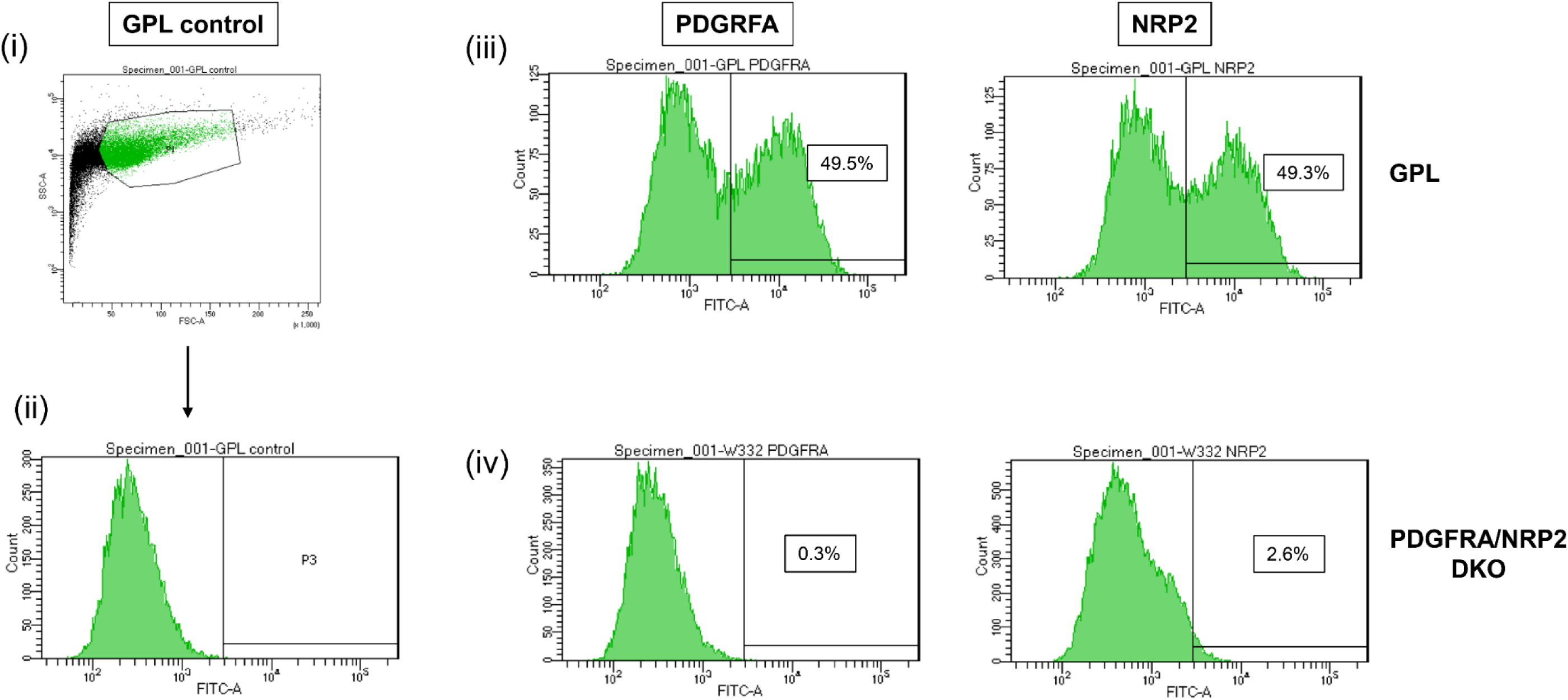
FACS analysis of GPL and NRP2/PDGFRA DKO cell lines. (i) Unlabeled GPL cells were gated (P1) in forward (FSC) and side-scatter (SSC) then shown as histogram (ii) with P3 gated for fluorescence. (iii) GPL cells labeled with PDGFRA or NRP2 antibodies for expression followed by donkey anti-goat FITC secondary antibody. Percent positive shown in histogram. (iv) PDGRFA/NRP2 DKO (W332) cells were similarly labeled with PDGFRA or NRP2 antibodies followed anti-goat FITC secondary antibody. Percent positive shown in histogram.

### Evaluation of guinea pig CD147 expression and interaction with GPCMV PC

A second cellular protein, CD147, was evaluated as a possible receptor for GPCMV PC based on the ability of human CD147 to influence HCMV endocytic infection. Additionally, CD147 is important for neurological and placental development (55, 56) and if a viral tropism factor to relvant tissues could contribute to the stigmata of cCMV. The guinea pig CD147 homolog exhibits 59% identity to human CD147 protein based on BLAST analysis of human and guinea pig predicted protein amino acid sequences. A codon optimized guinea pig CD147 synthetic gene was generated (Genscript) encoding a C-terminal FLAG epitope tag and cloned into pcDNA3.1(+) under HCMV IE enhancer control for transient plasmid expression studies (pDNAgpCD147). A protein of approximately 60 kDa was detected by anti-FLAG western blot of plasmid transfected GPL cell lysate, which was similar to the predicted size (Fig 5(i)). A cross-reacting antibody to CD147 (Antibodies Online) also detected a similar sized protein in GPL cells or plasmid transfected cells (Fig 5(ii)) indicating CD147 protein was endogenously expressed in GPL cells. Additionally, gpCD147 protein was the same size as endogenous CD147 identified by western blot in human MRC5 cell lysate (Fig 5(iv)). Guinea pig cell lines tested for CD147 expression included: GPASE; TEPI; REPI; GPL; and GPUVEC cells. Western blot analysis for CD147 expression identified a protein of approximately 60 kDa on all cell lysates tested (Fig 5(vi)). Levels of CD147 expression varied between cells lines based on lane loading control (β-actin) but was robustly expressed in GPL cells (Fig 5(vi)). Although CD147 was weakly expressed on GPASE cells, it was highly expressed on TEPI and GPUVEC cell lines (Fig 5) which are highly relevant cell types for cCMV.

**Figure 5.**
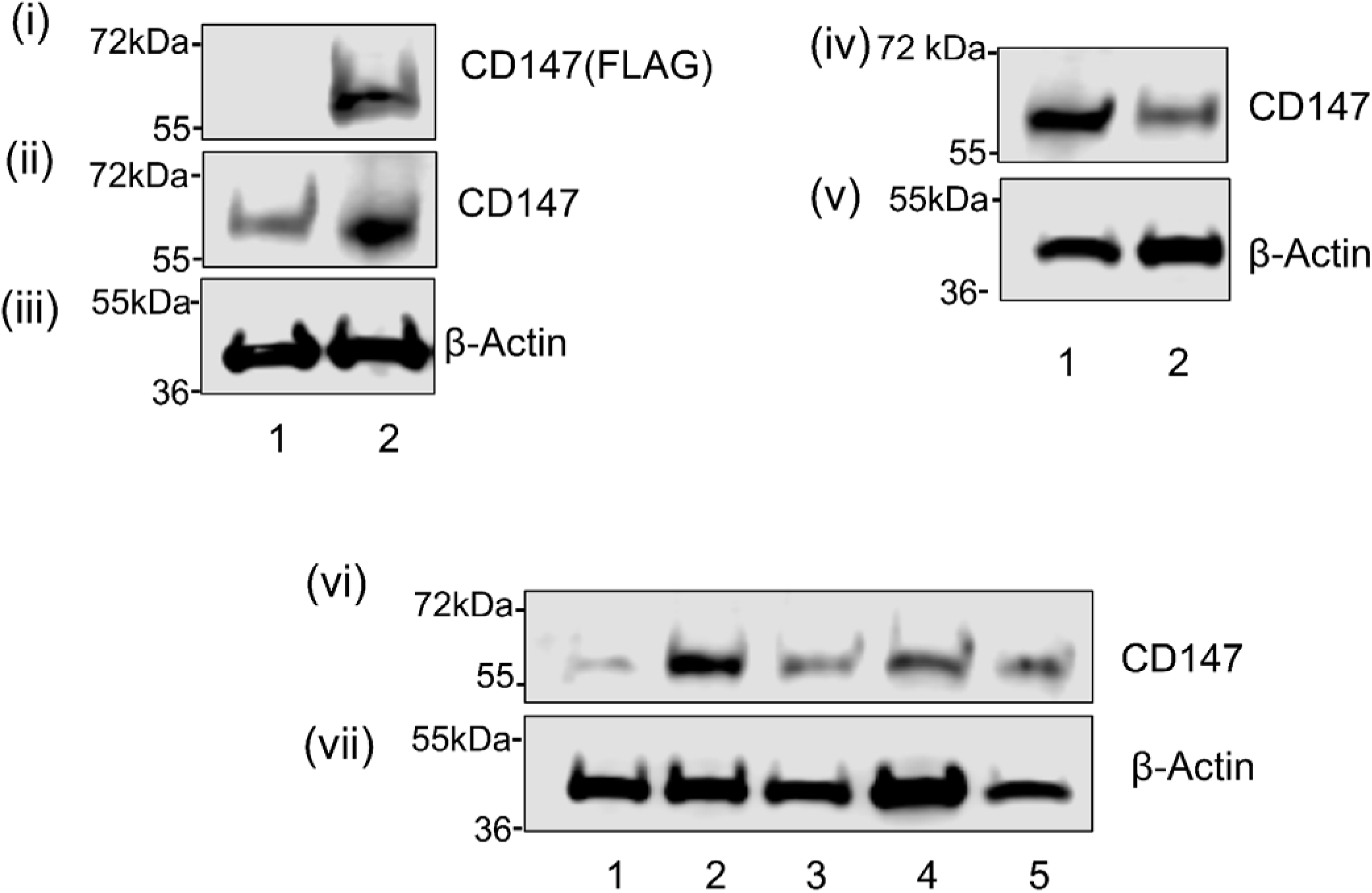
Detection of CD147 protein expression in guinea pig cells by western blot assay. (i) Western blot of transient plasmid FLAG-tagged CD147 synthetic gene protein expression on GPL cells. Lanes: 1, Control GPL cells; 2, CD147 plasmid. (ii) Western blot of plasmid and endogenous CD147 with anti-CD147 antibody. (iii) β-actin western blot lane loading control for (i & ii). (iv) Comparative endogenous CD147 expression in human MRC5 fibroblasts and GPL cells with anti-CD147 antibody. Lanes: 1, MRC5; 2, GPL cells. (v) β-actin western lane loading control for (iv). (vi) Western blot of endogenous CD147 expression on various guinea pig cell lines with anti-CD147 antibody. Lanes: 1, GPASE; 2, TEPI; 3, REPI; 4, GPL; 5, GPUVEC. (vii) β-actin western lane loading control for (vi). Protein detected with appropriate secondary antibody-HRP conjugate as described in materials and methods.

Subsequently, interaction of endogenous CD147 with GPCMV PC was evaluated by transient cellular expression of PC. Western blot analysis of RFP-trap IP assay demonstrated successful pulldown and interaction of individual components of the PC but failed to demonstrate CD147/PC interaction with CD147 only detected in the unbound flow through sample (Figure 6(i)). Evaluation of transient plasmid CD147 expression with PC also failed to demonstrate any interaction (data not shown). Interaction of CD147 with viral trimer (gH/gL/gO) was also evaluated by RFP-trap IP assay utilizing trimer components expressed by Ad vectors encoding individual trimer components (gH-GFP, gL-mcherry, and gO-FLAG) (24). Western blot of gH/gL/gO trimer IP assay successfully detected trimer formation but failed to demonstrate interaction with endogenous CD147 or NRP2 but did detect interaction with endogenous PDGFRA (Fig 6(ii)). We concluded that CD147 was not directly involved with GPCMV PC or trimer complex interaction and additionally that NRP2 did not interact with the viral trimer.

**Figure 6.**
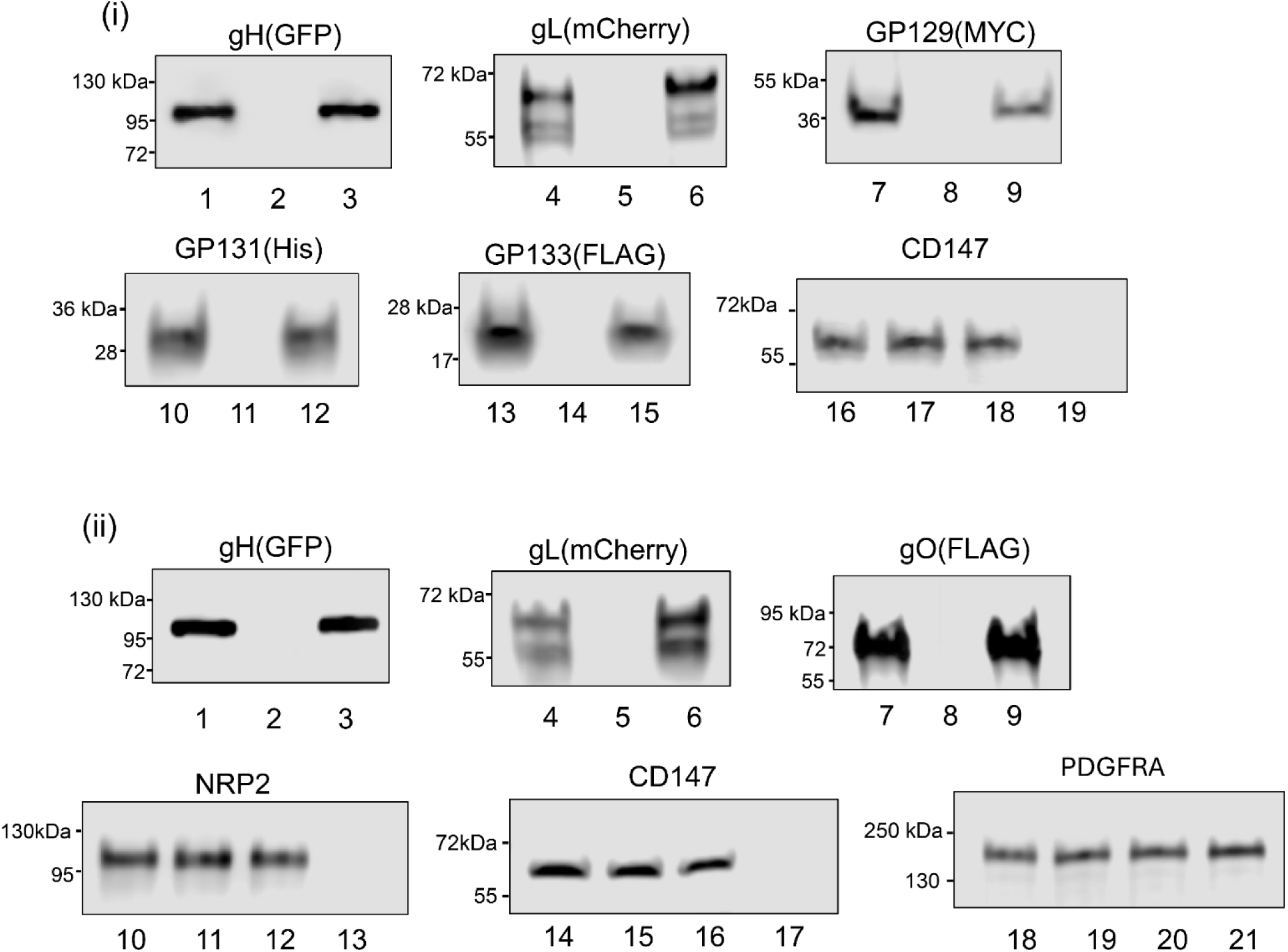
Immunoprecipitation (IP) of GPCMV PC or trimer with endogenous CD147 receptor. (i) Western blot of PC/CD147 IP assay. GPL cells transduced with all five recombinant Ad vectors encoding individual PC components (10 TDU/virus/cell). Samples analyzed as total cell lysate or processed for immunoprecipitation (IP) assay. IP carried out with RFP trap as previously described (25). IP lysate analyzed by western blot using respective primary antibodies for individual PC components or endogenous CD147: gH (α-GFP); gL (α-mCherry); GP129 (α-MYC); GP131 (α-His); GP133 (α-FLAG); endogenous CD147 (α-CD147). Protein detection visualized with appropriate secondary antibody-HRP conjugate as described in materials and methods. Western blot lanes: 1-3 (GFP); 4-6 (mCherry); 7-9 (MYC); 10-12 (His); 13-15 GP133 (FLAG); 16-19 CD147 (anti-CD147). Lanes 1, 4, 7, 10, 13, 16 (total cell lysate). Lanes 3, 6, 9, 12, 15,19 (IP). Lanes 2, 5, 8, 11, 14, 17 (mock control lysate). Lane 18 IP flow through. (ii) Control GPCMV trimer/receptor IP assay. Western blot of GPCMV gH/gL/gO trimer IP assay with endogenous candidate receptors (CD147, NRP2 and PDGFRA). GPL cells transduced with Ad vectors encoding individual trimer components (10 TDU/virus/cell) for interaction with endogenously expressed receptor proteins. IP lysate analyzed by western blot using respective primary antibodies for individual trimer components or endogenous receptor: gH (α-GFP); gL (α-mCherry); gO (α-FLAG); endogenous NRP2 (α-NRP2); CD147 (α-CD147); PDGFRA (α-PDGFRA). Western blot lanes: 1-3 (GFP); 4-6 (mCherry); 7-9 (FLAG); 10-13 (NRP2); 14-17 (CD147); 18-21 (PDGFRA). Lanes 1, 4, 7, 10, 14, 18 (total cell lysate). Lanes 3, 6, 9, 13, 17, 21 IP). Lanes 2, 5, 8, 11, 15, 19 (mock control lysate). Lanes 12, 16 and 20 IP flow through.

### Double knockout of PDGFRA and CD147 and impact on GPCMV infection

Next, CRISPR/Cas 9 strategy was employed to generate a double knockout of PDGFRA and CD147 on GPL cells with gRNAs targeting each gene following previously described protocol (25). A double knockout GPL cell line (designated CD147/PDGFRA DKO) was evaluated for ablation of target receptor protein expression by western blot of cell lysates from DKO and wild type GPL cells, which demonstrated the KO of PDGFRA and CD147 but NRP2 expression was retained (Fig 7(i-iii)). Specific alteration of PDGFRA and CD147 targeted exon gene sequence was also confirmed by PCR sequencing (data not shown). A comparative growth curve of GPCMV(PC+) on confluent monolayers in six well plates of GPL or DKO cells (MOI 1 pfu/cell) demonstrated that the DKO cells retained the ability to be infected by GPCMV (Fig 7(v)). However, GPCMV growth kinetics were attenuated with complete lysis of the cell monolayer occurring by 12 days post infection (dpi) compared to 6 dpi for normal GPL cells. Additionally, peak viral titer was approximately 3 logs lower for DKO cells compared to wild type GPL cells. Ectopic expression of CD147/PDGFRA on DKO cell line restored virus growth to near wild type level (Fig 7(vi)). In a separate set of experiments, endogenous surface expression of receptor proteins on GPL and CD147/PDGFRA DKO cells were compared by flow cytometry (Fig 8). This demonstrated the lack of PDGFRA protein on the surface of the DKO cells compared to GPL cells, which was predicted by western blot result (see Fig 7 and Fig 8). Evaluation of NRP2 endogenous protein surface expression on both GPL and DKO cells demonstrated comparative levels of NRP2 protein expression (Fig 8). CD147 surface expression was unable to be evaluated by flow cytometry because of a lack of available suitable antibody but based on western blot data (Fig 7) results should be similar to PDGFRA on the DKO cell line. We concluded that although CD147 did not interact directly with viral PC, it had the capacity to modify endocytic virus infection by an undetermined indirect method. A control comparative evaluation of HSV-1 infection (MOI 1 pfu/cell) on GPL and various receptor KO GPL cell lines was carried out (Fig 9) and demonstrated that HSV-1 infection was unaffected by single (PDGFRA) or double (NRP2/PDGFRA or CD147/PDGFRA) receptor knockout and that knockout of these proteins specifically impacted GPCMV infection (Fig 9).

**Figure 7.**
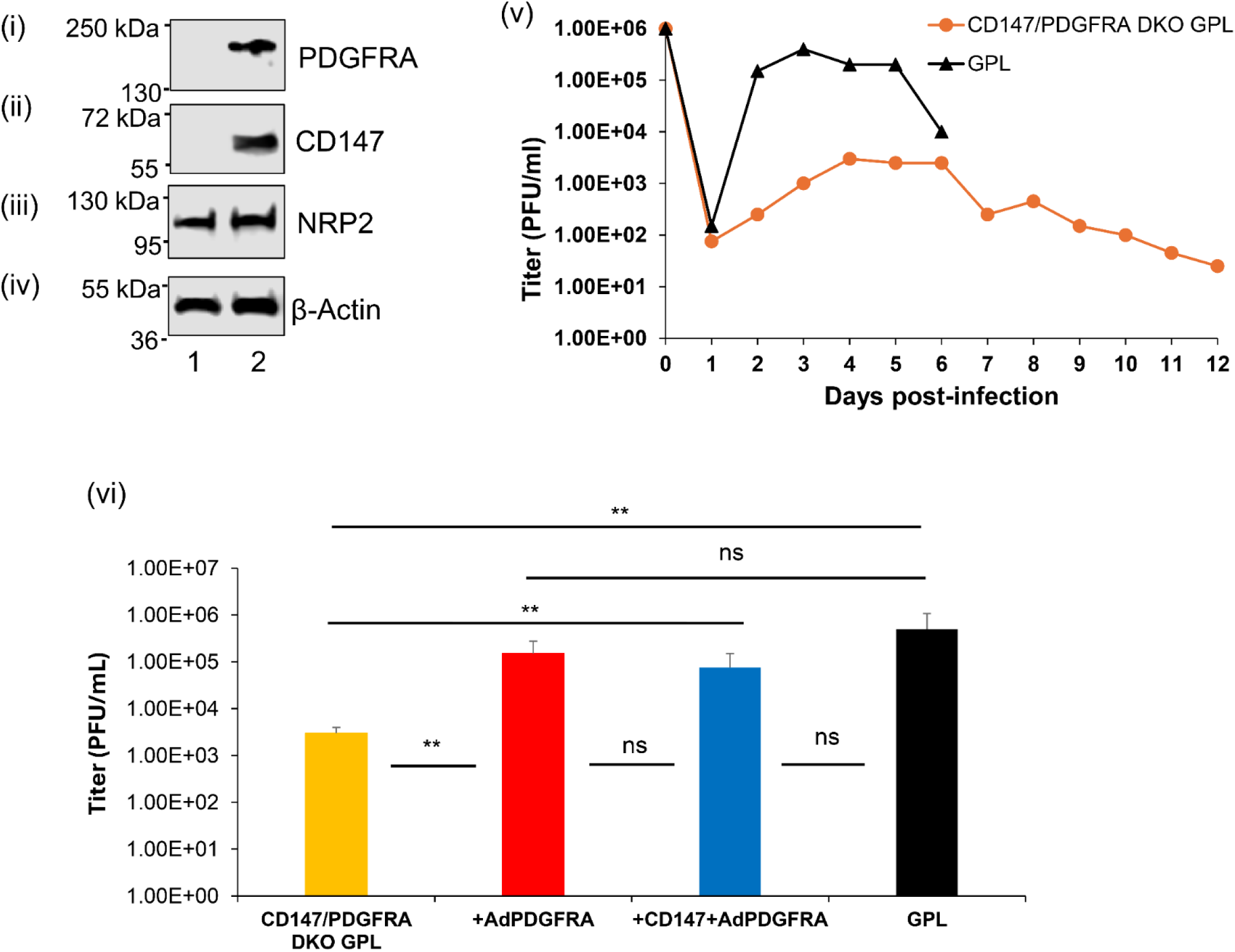
Characterization of CD147/PDGFRA DKO cells and evaluation of GPCMV infection. (i)-(iv) Receptor western blot of cell lysate. (i) Western blot for PDFGRA. (ii) Western blot for CD147. (iii) Western blot for NRP2. (iv) Western blot for lane loading control β-actin. Lanes: 1, DKO cells; 2, GPL cells. (v) Comparative growth curve of GPCMV(PC+) on GPL (black triangle) and DKO cells (orange circle). Virus infection MOI 1 pfu/cell with evaluation days 1-6 post infection (GPL) and days 1-12 (DKO). Results shown are average viral titer for experiment carried out in triplicate on 6 well plates with infection on confluent cell monolayers. (vi) Comparative GPCMV(PC+) growth (MOI 1 pfu/cell) on DKO (yellow) or GPL (black) cells or DKO cells ectopically expressing receptors PDGFRA (red) or CD147+PDGFRA (blue). Results shown are average virus titers at 3 days post infection from experiments performed in triplicate (\*\**p* < 0.01; ns = not significant).

**Figure 8.**
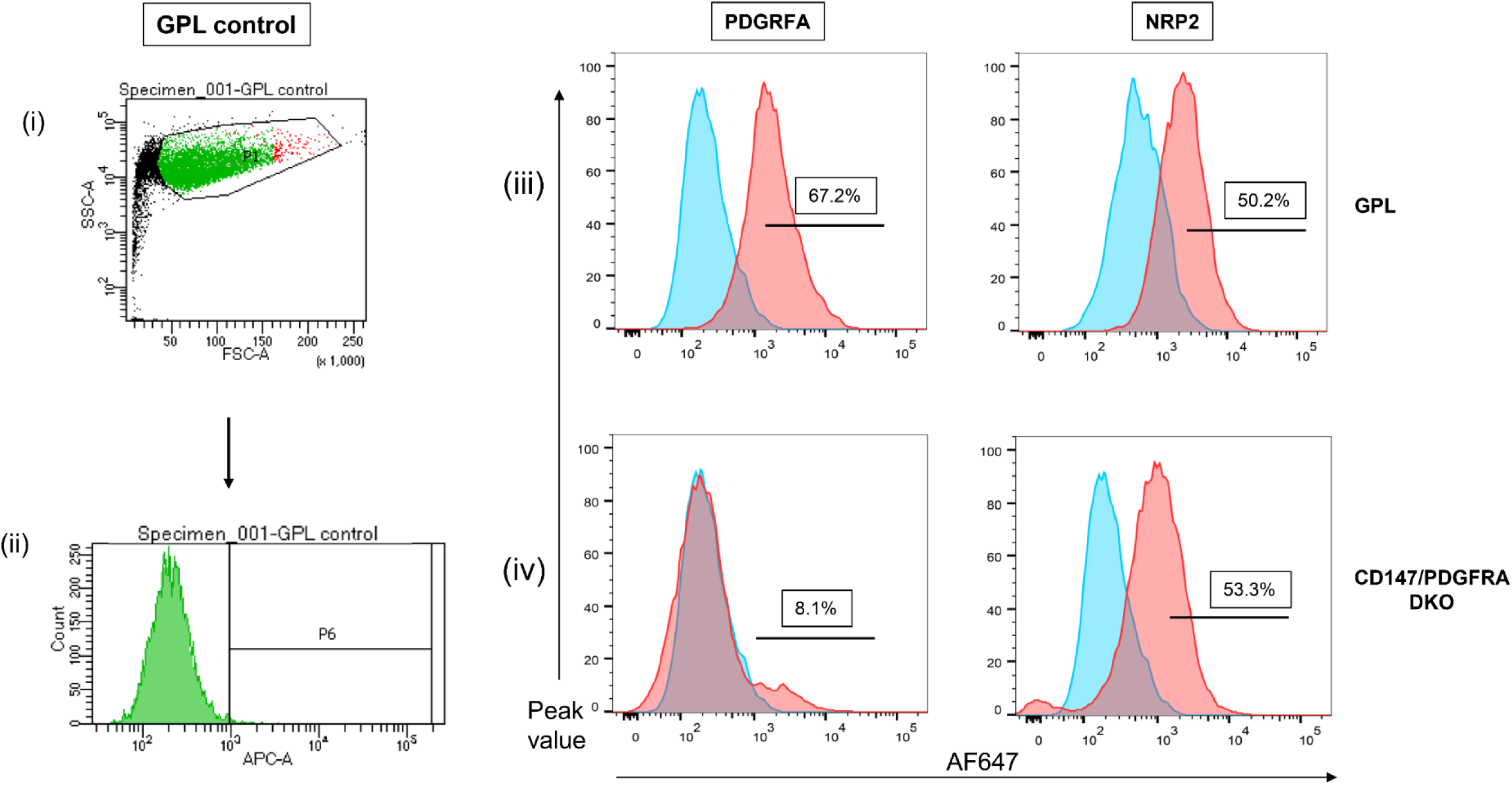
FACS analysis of GPL and CD147/PDGFRA DKO cell lines. (i) Unlabeled GPL cells were gated (P1) in forward (FSC) and side-scatter (SSC) then shown as histogram (ii) with P6 gated for fluorescence. (iii) GPL cells labeled with PDGFRA or NRP2 antibodies for expression followed by donkey anti-goat AF647 secondary antibody. Unlabeled GPL negative peak (blue) was overlayed with antibody labeled peak (red) to show comparison. Percent positive shown in histogram. (iv) CD147/PDGRFA DKO cells were similarly labeled with PDGFRA or NRP2 antibodies followed anti-goat AF647secondary antibody. Unlabeled GPL negative peak (blue) was overlayed with antibody labeled peak (red) to show comparison. Percent positive shown in histogram.

**Figure 9.**
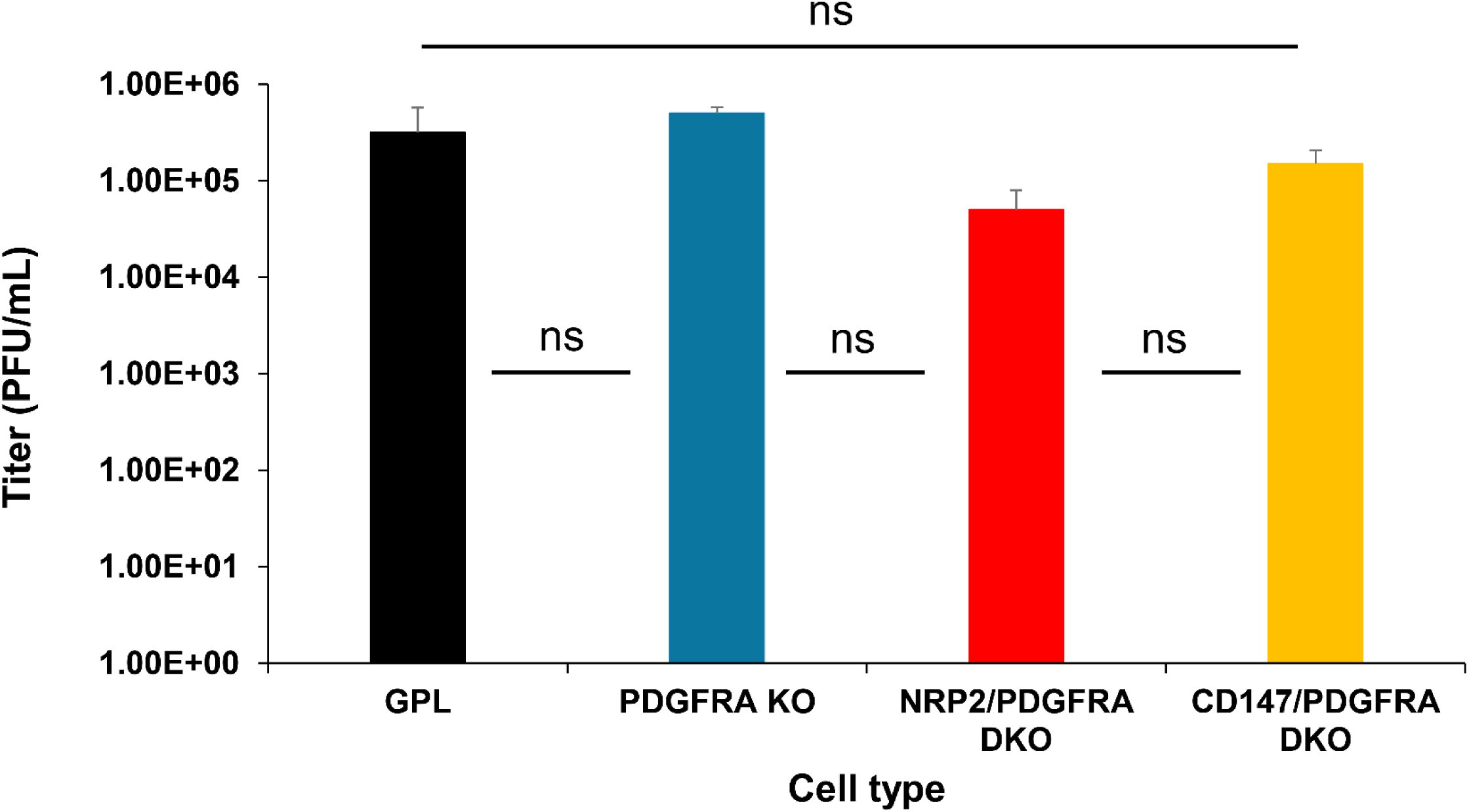
Evaluation of control virus HSV-1 infection on GPL and various KO cell lines. GPL or KO cell lines were infected with HSV-1 (MOI = 1 PFU/cell). Infected cell lysate harvested at 60 hpi and virus titered for each infected cell line. Results shown are average values for experiments carried out in triplicate. Infected cells: GPL (black); PGFRA KO (blue); NRP2/PDGFRA DKO (red); CD147/PDGFRA DKO (yellow). Statistical analysis was performed with student t-test (ns = not significant).

## Discussion

This study attempted to prevent GPCMV infection by alternative pathways of viral entry by targeted knockout of key candidate receptors in the only guinea pig cell line demonstrated to enable GPCMV infection by both direct and endocytic pathways via endogenously expressed cell receptors. Although GPCMV gB is the fusogenic protein essential for infection of all cell types and both pathways of entry, it is the ability of the gH/gL-based viral complexes to interact with key cellular receptors that dictate in part the pathway of cell entry. Additionally, the level of specific receptor protein expression on specific cell types might also influence a specific pathway of entry. Copy number of specific viral glycoprotein complexes on viral particles and affinity for specific receptors may also have a role to play in the pathway of entry (57, 58). Viral gHgLgO trimer interaction with PDGFRA enables direct cell entry in both HCMV and GPCMV but requires species specific receptor (28). Affinity for PDGFRA receptor may vary between HCMV strains due to high variability in the gO sequence, resulting in either reduced or enhanced binding for PDGFRA and subsequent entry via the direct pathway (59). GPCMV strains vary in predicted gO amino acid sequence with 22122 strain, which preferentially grows on fibroblast cells exhibiting 25% variability with clinical TAMYC strain gO where TAMYC strain preferentially infects non-fibroblast cell lines (i.e. PDGFRA negative cells) (60).

GPCMV PC enables virus infection via the endocytic pathway similar to HCMV. Understanding the interaction between CMV PC and receptor(s) is crucial for the development of antiviral therapies, insights into viral tropism, disease, and the development of new vaccine strategies. GPCMV PC and candidate receptor studies are additionally important to improve the translational impact of cCMV vaccine research in this model. Prior studies have determined the importance of the PC for GPCMV entry into various established guinea pig cell lines including epithelial, endothelial and trophoblasts, which lack PDGFRA receptor for direct cell entry (25, 29, 30, 51). Consequently, the PC is important for GPCMV dissemination and pathogenicity in the guinea pig including congenital infection (25, 30, 51) and likely required to target monocytes and establish viral latency (52). In HCMV, varying levels of PDGFRA on placental trophoblasts results in different pathways of cell entry and this is also possibly dictated by the ratio of trimer to PC on virion particle (33). An established guinea pig trophoblast (TEPI) cell line lacks PDGFRA with GPCMV infection via the endocytic pathway (51). However, ectopic expression of PDGFRA on TEPI cells enables infection via direct pathway independent of the PC (28). Vaccine strategies encoding the PC can enhance protection against cCMV in the guinea pig demonstrating the importance of the PC as a neutralizing target antigen along with gB against GPCMV (26, 61).

In HCMV, PC-based endocytic cell entry is only partially understood and poorly defined in animal CMV models. Consequently, this study sought to evaluate candidate receptors for PC-based GPCMV cell entry in order to gain a better understanding of GPCMV infection and investigate the similarity between HCMV and GPCMV infection. Additionally, we sought to determine if it was possible to block both pathways of viral cell entry by elimination of key receptors. Candidate PC receptors were selected based on best defined for HCMV (NRP2), or potential for impact related to enhanced virus tropism/infection especially for placental or neurological infection based on candidate receptor abundance (CD147). Based on the human protein atlas (proteinatlas.org), both NRP2 and CD147 are widely expressed in the human body including the placenta. The guinea pig homolog proteins were highly expressed in all guinea pig cell lines including fibroblast, epithelial and endothelial cell types and especially in those cells highly relevant to cCMV. Perhaps NRP2 is the most studied PC receptor for HCMV of epithelial and endothelial cells (45, 49, 50). In this report, we demonstrate that the guinea pig homolog NRP2 protein functions as a receptor for GPCMV PC and is widely expressed in various guinea pig cell lines. Consequently, it is likely a PC receptor for GPCMV entry into various cell type via endocytic pathway. However, it is not assumed that NRP2 is uniquely able to directly interact with PC for virus infection and this may vary with specific cell type, abundance of receptor and affinity for the PC. Indeed, ThBD has been suggested to be a more favorable HCMV PC receptor in conjunction with gB (49). A general limitation of guinea pig studies is the availability of reagents. The lack of an available cross reacting ThBD antibody prevented additional studies on guinea pig ThBd receptor candidate at present time.

Although various HCMV PC receptor candidates have been identified for HCMV, only one receptor (PDGFRA) is demonstrated to interact with the trimer complex to enable direct cell entry independent of the PC (62, 63). Guinea pig PDGFRA was previously demonstrated to enable direct entry and knockout of this receptor blocked direct but not PC-dependent endocytic cell entry in GPL cells (26). Consequently, complete prevention of GPCMV cell infection would require the absence of both direct pathway receptor (PDGFRA) and PC endocytic entry pathway receptor(s). If infection is successfully blocked by knockout of both PDGFRA (direct entry) and an additional receptor, then this receptor is likely at least involved in, if not responsible for, PC endocytic entry. Guinea pig NRP2 interacted with PC and protein was expressed on the cell surface and in multiple cell lines capable of endocytic infection suggesting that it likely acts as a viral PC receptor. Knockout of both PDGFRA and NRP2 resulted in full inhibition of GPCMV infection without impacting infection of control HSV-1 on the NRP2/PDGFRA DKO cell line. This demonstrated that inhibition as a result of receptor knockout was specific for GPCMV. Potentially, this establishes the foundation for the development of receptor-specific small molecule or peptide-based antivirals, enabling evaluation of their protective efficacy against cCMV in this model (64). Since NRP2 is also used by other viruses as an entry receptor, including Lujo and rabies virus, this approach may give rise to a more broad-spectrum antiviral strategy (65, 66). Likely, soluble NRP2 recombinant protein could be used to evaluate inhibition of endocytic infection on other guinea pig cell types but this was considered outside the scope of this initial report. It was important to demonstrate that GPCMV utilizes guinea pig NRP2 as an endocytic entry receptor in association with PC as this extends the similarity to HCMV but in a species-specific manner. In contrast to HCMV, murine CMV (MCMV) utilizes NRP1 as a general cell entry receptor and importantly lacks a PC where elimination of NRP1 impacted infection of endothelial, fibroblast and macrophage cells (67). Similarly, NRP1 is also identified as an entry receptor for Kaposi’s sarcoma-associated herpesvirus (KSHV) (68) which also lacks a homolog PC. In future studies it would be important to evaluate receptors on other cell types but based on HCMV research there are likely to be additional multiple candidate receptors that should be tested.

CD147 (also known as BSG or EMMPRIN) is a single-chain type I transmembrane protein with two immunoglobulin-like domains in its extracellular region (69). CD147 is highly expressed in the placenta based on the Human Protein Atlas and was evaluated as a possible receptor for GPCMV PC based on the ability of human CD147 to influence HCMV endocytic infection (47). CD147 is important for neurological and placental development (55, 56) with the potential for cCMV disease based on this receptor as a viral tropism factor for specific tissues. This candidate receptor was detected on all guinea pig cell lines, including placental trophoblast, amniotic sac cells and GPUVEC cells supporting its further evaluation. Similar to HCMV, GPCMV PC did not interact with CD147 in IP assays but knockout of CD147 did specifically decrease GPCMV infection without affecting HSV-1 growth. In HCMV, a CD147 shRNA knockdown strategy in endothelial cells resulted in reduced HCMV infection but this strategy was unsuccessful in epithelial cells because of reported toxicity to the cells by shRNA expression (47). Human dermal fibroblast cells also used in parallel HCMV studies expressed between 5-10% the level of CD147 protein compared to human endothelial and epithelial cells. Perhaps given the low level of endogenous expression, CD147 shRNA knockdown in dermal fibroblasts had no significant impact on HCMV infection (47). It remains to be determined if there is an impact on other more commonly studied human fibroblast cells lines (eg. MRC5) where CD147 protein is more highly expressed (Fig 5). Additionally, in the study by Johnson and colleagues, overexpression of CD147 enhanced virus entry into HeLa cells (47). In contrast to dermal fibroblasts in HCMV study, the GPL fibroblasts used in the current studies expressed robust levels of CD147, NRP2 and PDGFRA. Overexpression of CD147 or NRP2 on GPL cells did not enhance infection, except in the backdrop of specific KO. Potentially, CD147 can modify levels of cellular integrins on the cell surface and integrins are considered a potential co-receptor for HCMV infection by interaction with gB (70–72). Therefore, modulation of integrin protein expression might impact infection of GPCMV in a similar manner to HCMV (56, 73). However, the lack of available antibodies to enable evaluation of various guinea pig integrins expressed on the cell surface in the presence or absence of CD147 prevented any evaluation of this aspect for GPCMV infection at present time.

Since endogenous CD147 failed to be detected in PC IP assays in contrast to NRP2, it seems unlikely that NRP2 and CD147 directly interact on the cell surface. Although, it might be possible that knockout of CD147 indirectly impacts on the surface expression of NRP2 this was not demonstrated by flow cytometry assays (Fig 8). The PC enables cell free virus infection by endocytic pathway, it also allows viral cell-to-cell spread in the absence of cell free virus. This was demonstrated for GPCMV in studies with a gO knockout mutant in the backdrop of a GPCMV (PC+) virus with cell-to-cell spread across both fibroblast and epithelial cell monolayers (25). In contrast, a gO knockout mutant virus in the backdrop of PC-negative virus is a lethal mutation (24, 25). Potentially, cell-to-cell spread enables the virus to evade neutralizing antibodies targeting cell-free virus. Presumably, similar PC receptors enable cell-to-cell spread but this remains to be evaluated for both HCMV and GPCMV and worthy of future studies but outside the scope of this initial report.

In summary, this study highlights the importance of cellular receptors PDGFRA and NRP2 in facilitating GPCMV infection. PDGFRA interacts with the viral trimer to mediate direct entry while NRP2 interacts with PC to promote infection via endocytic pathway. However, based on HCMV studies, it is unlikely that NRP2 is the only candidate receptor for GPCMV PC endocytic cell entry. The study of endocytic CMV infection is additionally complicated by the ability of specific cellular proteins to indirectly modulate endocytic viral infection (eg. CD147). Future efforts to better understand GPCMV PC-dependent infection should evaluate infection of non-fibroblast cells especially those most relevant to cCMV. These were considered outside the scope of this initial report and a limitation to current findings. Overall, results demonstrate that GPCMV exhibits conservation with HCMV for key receptors (PDGFRA and NRP2) and cell entry pathways as well as indirect effect by CD147. This contrasts with murine CMV, which lacks a PC and utilizes different cell receptors with infection perhaps more similar to KSHV and EBV. Given the significance of the PC as a key vaccine target, these findings continue to emphasize the translational importance of PC-based vaccine studies in this CMV small animal model.

## CRediT authorship contribution statement

K. Yeon Choi: Conceptualization, Methodology, Investigation, Data curation, Formal analysis, Writing - original draft, Writing - review & editing. Yushu Qin: Methodology, Investigation, Data curation, Formal analysis, Writing – original draft, Writing - review & editing. Nadia S. El-Hamdi: Conceptualization, Methodology, Investigation, Data curation, Formal analysis, Writing – original draft, Writing - review & editing. Alistair McGregor: Conceptualization, Methodology, Investigation, Data curation, Formal analysis, Writing - original draft, Writing - review & editing, Funding acquisition.

## Acknowledgements

The authors would like to thank Austin Selman for his technical assistance in some tissue culture experiments. We are grateful to M. A. McVoy (VCU) for the generous gift of the second-generation GPCMV BAC. This study was supported by grants from National Institute of Health (NIH) institutes NIAID and NICHD, awarded to AM: R01AI100933; R01AI098984; R01HD090065.

